# Diverse paths for chemoreception in ciliated neurons contacting the cerebrospinal fluid in the spinal cord

**DOI:** 10.64898/2026.04.14.718607

**Authors:** Emily Verran, Louise Moizan, Loeva Tocquer, Feng B. Quan, Claire Wyart

## Abstract

Cerebrospinal fluid-contacting neurons (CSF-cNs) are mechanosensory cells in the spinal cord that detect compression and regulate locomotion, posture, and morphogenesis. Although CSF-cNs respond to changes in pH, neurotransmitters and metabolites, their chemosensory repertoire is not fully understood. Using hybridization chain reaction, we investigated the distribution of expression of chemoreceptors in CSF-cNs and neighboring cells in the spinal cord. We found that CSF-cNs express receptors for glutamate (*grm2*), somatostatin (*sstr2*) and low-density lipoprotein (LDL) (ldlrad2), indicating roles in detecting glutamate, somatostatin and LDL in the CSF. High LDL receptor expression in CSF-contacting cells suggests CSF lipid capture. Most receptors were enriched but not exclusive to CSF-cNs and also appeared in ependymal radial glial cells. Our findings indicate multiple chemosensory pathways can sustain long-distance communication between neurons and glia through the cerebrospinal fluid.

## INTRODUCTION

Interoception is a major path through which the nervous system assays internal states (Wyart et al., 2023; Verdonk et al., 2025). The cerebrospinal fluid (CSF) is a complex solution in which the central nervous system baths. The CSF is produced by the choroid plexus and changes composition as a function of age, time of the day, and physiological conditions (Parnetti et al., 2019; Fagan et al., 2021; Hermann et al., 2021) so that spinal taps are often used to find out about the nature of an infection or a neurological disease. Cells in contact with the cerebrospinal fluid include ependymal cells, radial glia, tanycytes, secretory cells from the choroid plexi, the subcommissural organ or the pineal gland as well as ciliated neurons. In the last decade, a sensory pathway involving ciliated neurons detecting changes of the cerebrospinal fluid was discovered as important to adjust posture during challenging behaviors, locomotor speed, as well as morphogenesis and even innate immunity (Wyart et al., 2023). A century ago, Kolmer and Agduhr described ciliated neurons at the level of the central canal in the spinal cord of over a hundred vertebrate species that contact the cerebrospinal fluid (CSF) (Kolmer, 1921; Agduhr, 1922; Dale et al., 1987). These CSF-contacting neurons (CSF-cNs) are dense GABAergic neurons in the spinal cord of macaques, mice, turtles or zebrafish (Djenoune and Wyart, 2017).

During development, CSF-cNs are organised in two rows originating from different progenitor domains, pMN and p3 in zebrafish (Shin et al., 2007; Yang et al., 2010; Huang et al., 2012; Yang et al., 2020) versus p3 and p2 in mice (Petracca et al., 2016; Di Bella et al., 2019). Early during embryonic development, a bidirectional flow of CSF in the central canal can enable the effective transport of large particles (Thouvenin et al., 2020). Later in the central canal of mature animals, the CSF typically flows from anterior to posterior and its flow can be enhanced by muscle contractions (Thouvenin et al., 2020). By interacting with the Reissner fiber (Reissner, 1860), a polymer under tension in the CSF, CSF-cNs detect spinal compression (Bohm et al., 2016; Jalalvand et al., 2016b; Orts-Del’Immagine et al., 2020). This process requires the channel TRPP2 or PKD2L1 that is a highly specific marker of CSF-cNs across species (Sternberg et al., 2018; Djenoune et al., 2014; Petracca et al., 2016; Gerstmann et al., 2022; Nakamura et al., 2023). In return, CSF-cNs acutely modulate locomotion, its speed and fine postural control (Wyart et al., 2009; Fidelin et al., 2015; Hubbard et al., 2016; Wu et al., 2021). On longer time scales, these cells, together with the Reissner fiber, control morphogenesis (Sternberg et al., 2018; Cantaut-Belarif et al., 2018; Troutwine et al., 2020; Rose et al., 2020; Marie-Hardy et al., 2023) via release of peptides from the Urotensin 2 family (Zhang et al., 2018; Bearce et al., 2022; Gaillard et al., 2023). In addition to urotensin related peptides, CSF-cNs are GABAergic neurons strikingly synthesizing numerous monoamines and peptides that differ between species: somatostatin in fish and lamprey, dopamine in lamprey, serotonin transiently in fish (Djenoune et al., 2017), trace amines in rodents (Hökfelt et al., 1973; Jaeger et al., 1983; Ren et al., 2017), urotensin related peptides in fish (Quan et al., 2015) among many others (Djenoune and Wyart, 2017).

The chemosensory functions of CSF-cNs are however the first ones that have been investigated. Over 20 years ago, Stoeckel et al. showed that these cells express specific P2X receptors for ATP (Stoeckel et al., 2003). Zuker’s team found that CSF-cNs firing is modulated by pH (Huang et al., 2006), which is later confirmed that the activity of CSF-cN is strongly modulated by changes in pH and osmolarity in mice and lamprey (Orts-Del’Immagine et al., 2012; Jalalvand et al., 2016a). CSF-cNs express numerous chemoreceptors and respond to bacterial metabolites upon invasion of the CSF by pathogenic bacteria triggering meningitis (Prendergast et al., 2023). Recent evidence for expression of opioid receptors in CSF-cNs indicate a role in spinal cord injury (Yue et al., 2024). Yet, the extent of their chemosensory abilities, i.e. which receptors CSF-cNs express and molecules they can detect in the CSF, is not fully understood.

To identify new pathways enabling long range signaling in the CSF via CSF-contacting neurons, we investigated here in the CSF-cN transcriptome in larval zebrafish which novel putative chemoreceptors may be expressed and enriched in these cells. To confirm expression in CSF-cNs and characterize the spatial expression pattern, we used hybridization chain reaction (HCR) on 2-3 days post fertilization (dpf) larvae and found that both dorsolateral and ventral CSF-cNs expressed the low-density lipoprotein (LDL) receptor 2 *ldlrad2*, glutamate receptor 2 *grm2a* and *ptprna*. In contrast, somatostatin receptor 2 *sstr2a* was specifically expressed in the ventral population while somatostatin itself is only present in the dorsal population. Interestingly, multiple of these receptors were also found in other cells contacting the CSF such as ependymal radial glia (ERGs) or floor plate neuroepithelial cells (FP). Altogether, our results indicate that numerous chemosensory signaling pathways can enable long distance communication at the CSF interface.

## RESULTS

### Selection of receptors combining enrichment in CSF-cNs and high expression

To identify novel chemoreceptors in CSF-cNs, we investigated the transcriptome of these cells performed out of 5 replicates run after sorting for GFP+ cells in the 3-day old guillotined double transgenic *Tg(pkd2l1:GAL4;UAS:GFP)* larvae (Prendergast et al., 2023). We selected receptors who were enriched (Log Fold Change > 1.15) and had an absolute number of reads in the GFP+ CSF-cNs population above 10 Fragments Per Kilobase per Million mapped fragments (FPKM) (**Table 1**). Four receptors caught our attention: the low-density lipoprotein (LDL) receptor 2 (*ldlrad2*), the somatostatin receptor 2a (*sstr2a*), the glutamate metabotropic receptor 2 (*grm2a*) and the *ptprn* receptor (*ptprna*).

**Table 1.** Selection of receptors highly expressed and enriched in CSF-cNs from a previous transcriptome (data extracted from Prendergast *et al*., 2023). Read counts in Fragments Per Kilobase per Million mapped fragments (FPKM) per transcripts were obtained from 5 replicates for both GFP+ and GFP-cells in pooled *Tg(pkd2l1:GAL4; UAS:GFP)* transgenic larvae.

| Description | Zv10 Feature | Mean FPKM GFP negative | Mean FPKM GFP positive | Log Fold Change | Probability | False Discovery Rate |
| --- | --- | --- | --- | --- | --- | --- |
| Low Density Lipoprotein Receptor Class A Domain Containing 2 | *ldlrad2* | 6,28 | 32,33 | 2,49 | 8,50. 10<sup>-10</sup> | 4,46. 10<sup>-7</sup> |
| Glutamate Receptor, metabotropic 2a | *grm2a* | 2,64 | 12,2 | 2,25 | 7,29. 10<sup>-7</sup> | 1,81. 10<sup>-4</sup> |
| Somatostatin Receptor 2a | *LOC100333578* | 7,89 | 20,76 | 1,42 | 1,00. 10<sup>-3</sup> | 4,95. 10<sup>-2</sup> |
| Protein Tyrosine Phosphatase Receptor Type Na | *ptprna* | 6,87 | 14,71 | 1,16 | 8,46. 10<sup>-4</sup> | 4,47. 10<sup>-2</sup> |

### Somatostatin receptor *sstr2a* is expressed in ventral CSF-cNs and unknown dorsal spinal cells in zebrafish larvae

To confirm the expression of *sstr2a* in CSF-cNs, we performed hybridization chain reaction (HCR) for *sstr2a* and *pkd2l1* transcripts on 3-day post fertilization (dpf) AB *mifta* ^*-/-*^ larvae (n = 9 fish, **Figure 1**). We used DAPI for delaminating the ventral and dorsal boundaries of the spinal cord and highlighting the central canal (**Figure 1A1**). The *pkd2l1* HCR probe nicely labels as shown before (Djenoune et al., 2014) both dorsolateral (triangle) and ventral (arrowhead) CSF-cNs (**Figure 1A2, A2i**). We observed all along the spinal cord co-expression of the receptor *sstr2a* with *pkd2l1+* cells located ventrally from the central canal, indicating that ventral CSF-cNs (arrowhead, **Figure 1A3, A3i, A4, A4i, C, D**) but not dorsolateral CSF-cNs (triangle, **Figure 1B3, B3i, B4, B4i, C, D**) express the *sstr2a* receptor. In addition, we noticed high expression of *sstr2a* is also found in unknown dorsal most cells on the lateral edge of the spinal cord (asterisk, **Figure 1B3, B4, C**). The total number of *pkd2l1*^+^ CSF-cNs was similar between ventral (18 ± 2.0 per 100 µm) and dorsolateral (18 ± 1.2 per 100 µm) CSF-cN populations. Quantification confirmed that *sstr2a* expression was largely confined to ventral CSF-cNs: on average, 12 ± 1.2 ventral *pkd2l1*^+^ CSF-cNs per 100 µm expressed *sstr2a*, compared to only 1 ± 0.1 dorsolateral cell. These results demonstrate that *sstr2a* expression is largely restricted to ventral CSF-cNs among the CSF-cN population.

**Figure 1.**
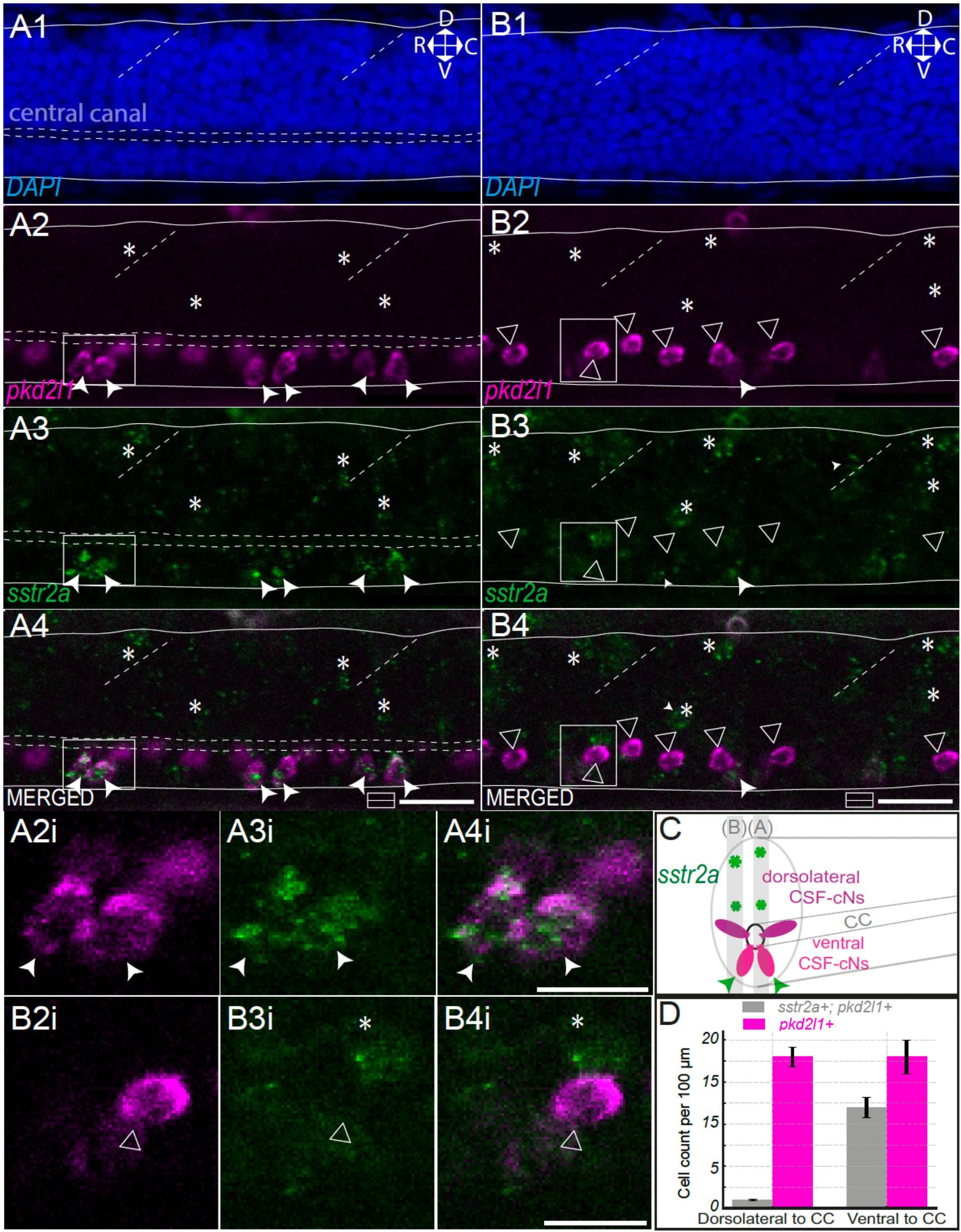
The somatostatin receptor *sstr2a* is predominantly expressed in ventral CSF-cNs and unidentified dorsal spinal cells. **(A)** Midline sagittal optical section of the spinal cord of a 3 dpf larval zebrafish showing DAPI staining **(A1)** outlining dorsal and ventral boundaries (solid lines), the central canal (horizontal dashed lines), and somite boundaries (oblique dashed lines). Orientation: dorsal (D), ventral (V), rostral (R), caudal (C). *pkd2l1* labeling (**A2**, magenta) marks dorsolateral (triangle) and ventral (arrowhead) CSF-contacting neurons (CSF-cNs). *sstr2a* expression (**A3**, green) is detected almost exclusively in ventral *pkd2l1*^+^ CSF-cNs and in a few unidentified dorsal spinal cells (asterisk). Scale bar, 20 µm. Insets (**A2–A4i)** show magnified ventral CSF-cNs. Scale bar, 10 µm. **(B)** Lateral sagittal section showing *sstr2a* expression (**B3**, green) in ventral *pkd2l1*^+^ CSF-cNs (B2, magenta, filled arrowhead) and in unidentified dorsal *pkd2l1*^−^ cells (asterisk). Dorsolateral *pkd2l1*^+^ CSF-cNs (triangle) lack *sstr2a* expression. Scale bar, 20 µm. Insets **(B2–B4i)** show a magnified dorsolateral *pkd2l1*^+^/*sstr2a*^−^ CSF-cN and a *sstr2a*^+^/*pkd2l1*^−^ dorsal cell (asterisk). Scale bar, 10 µm. **(C)** 3D schematic of a transverse spinal cord section summarizing *sstr2a* expression (green) primarily in ventral *pkd2l1*^+^ CSF-cNs (magenta) and in additional dorsal cells (asterisk). **(D)** Quantification of *sstr2a*^+^ CSF-cNs per larva (*n* = 3), normalized per 100 µm. In dorsolateral position to the central canal (CC): 1 ± 0.1 *sstr2a*^+^*/pkd2l1*^+^ cells per 100 µm (grey); in ventral position to the central canal (CC): 12 ± 1.2 *sstr2a*^+^*/pkd2l1*^+^ cells per 100 µm. Total *pkd2l1*^+^ cells (magenta): 18 ± 1.2 dorsolateral CSF-cNs per 100 µm and 18 ± 2.0 ventral CSF-cNs per 100 µm. Mean values are given ± s.e.m.

### Glutamate metabotropic receptor *grm2a* is expressed in ventral and dorsolateral CSF-cNs and additional dorsal spinal cells

To confirm the expression of *grm2a* in CSF-contacting neurons (CSF-cNs), we performed HCR for *grm2a* and *pkd2l1* transcripts on 2 dpf *AB mifta* ^*-/-*^ larvae (*n* = 6 fish; **Figure 2**). DAPI staining was used to delineate the ventral and dorsal boundaries of the spinal cord and to highlight the central canal (**Figure 2A1**). The *pkd2l1* HCR probes labeled both dorsolateral (triangle) and ventral (arrowhead) CSF-cNs (**Figure 2A2, B2**). Along the entire spinal cord, we observed co-expression of *grm2a* with all *pkd2l1*^+^ cells, indicating that both ventral (arrowhead; **Figure 2A3, A4, A4i, B3, B4, B4i**) and dorsolateral (triangle) CSF-cNs express the *grm2a* receptor. In addition, a few *grm2a*^+^ cells not expressing *pkd2l1* near the central canal were labeled (asterisk; **Figure 2B4, B4i**), as well as unidentified dorsalmost located spinal cells (asterisk; **Figure 2A3, A4, B1**). The total number of *pkd2l1*^+^ CSF-cNs was (19 ± 1.3 per 100 µm) for ventral and (14 ± 2.5 per 100 µm) for dorsolateral populations. Quantification confirmed that *grm2a* was expressed in both ventral and dorsolateral CSF-cNs (on average, 15 ± 1.7 per 100 µm for ventral CSF-cNs and 10 ± 2.8 per 100 µm for dorsolateral *pkd2l1*^+^ CSF-cNs per 100 µm were *grm2a*^+^).

**Figure 2.**
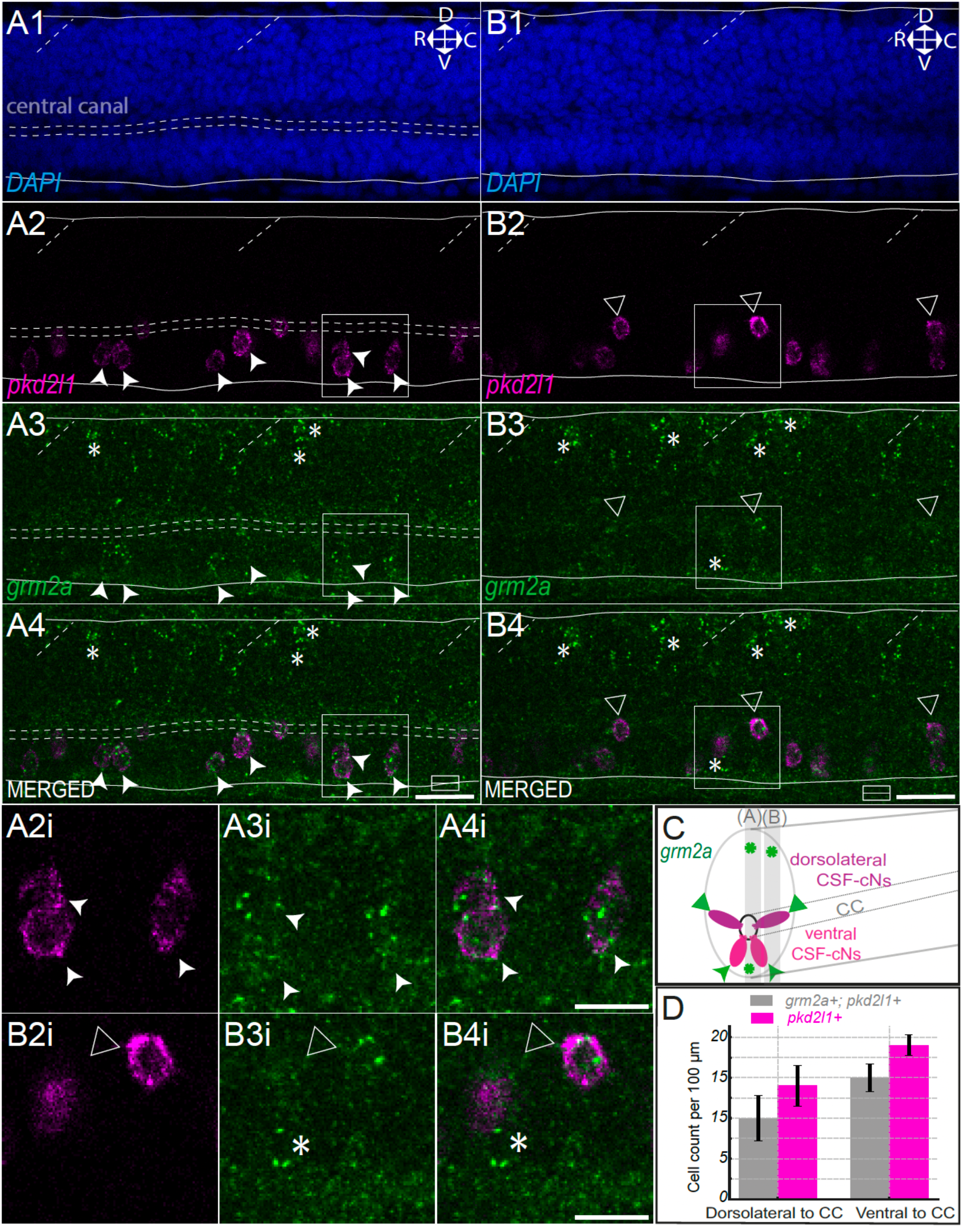
The metabotropic glutamate receptor 2 *grm2a* is expressed in ventral and dorsolateral CSF-cNs as well as in unknown dorsal cells in the spinal cord of larval zebrafish. **(A)** Sagittal optical section of the spinal cord of a 2 dpf larval zebrafish showing DAPI staining (A1) outlining dorsal and ventral boundaries (solid lines), the central canal (horizontal dashed lines), and somite boundaries (oblique dashed lines). Orientation: dorsal (D), ventral (V), rostral (R), caudal (C). *pkd2l1* labeling (A2, magenta) marks dorsolateral (triangle) and ventral (arrowhead) CSF-contacting neurons (CSF-cNs). *grm2a* expression (A3, green) overlaps with *pkd2l1* in both populations and is also detected in unidentified dorsal cells (asterisk). Scale bar, 20 µm. Insets (A2–A4i) show magnified ventral CSF-cNs. Scale bar, 10 µm. **(B)** Lateral sagittal section showing *grm2a* expression (**B3**, green) in dorsolateral *pkd2l1*^+^ CSF-cNs (**B2**, magenta, triangle) and in *pkd2l1*^−^ cells near the central canal and in unidentified dorsal cells (asterisk). Scale bar, 20 µm. Insets (**B2–B4i**) show a magnified dorsolateral CSF-cN (*pkd2l1*^+^, triangle) and a *grm2a*^+^/*pkd2l1*^−^ cell (asterisk). Scale bar, 10 µm. **(C)** 3D schematic of a transverse spinal cord section summarizing *grm2a* expression (green) in ventral and dorsolateral *pkd2l1*^+^ CSF-cNs (magenta), in additional dorsal cells and cells near the central canal (asterisk). **(D)** Quantification of *grm2a*^+^ CSF-cNs per larva (*n* = 3), normalized per 100 µm. In dorsolateral position to the central canal (CC): 10 ± 2.8 per 100 µm *grm2a*^+^/*pkd2l1*^+^ cells (grey); in ventral position to the central canal (CC): 15 ± 1.7 *grm2a*^+^/*pkd2l1*^+^ cells per 100 µm (grey). Total *pkd2l1*^+^ cells (magenta): 14 ± 2.5 dorsolateral CSF-cNs per 100 µm and 19 ± 1.3 ventral CSF-cNs per 100 µm. Mean values are given ± s.e.m.

### The phosphatase receptor *ptprna* is expressed predominantly in both ventral and dorsolateral CSF-cNs in 3 day old larval zebrafish

To confirm the expression of *ptprna* in CSF-cNs, we performed HCR for *ptprna* and *pkd2l1* transcripts on 3 dpf AB *mifta* ^-/-^ larvae (n = 5 fish, **Figure 3**). DAPI was used to delaminate the ventral and dorsal boundaries of the spinal cord and highlight the central canal (**Figure 3A1**). The *pkd2l1* HCR probe labels both dorsolateral (triangle) and ventral (arrowhead) CSF-cNs (**Figure 3A2, B2**). We observed all along the spinal cord co-expression of the receptor *ptprna* and *pkd2l1* in both ventral CSF-cNs (arrowhead, **Figure 3A3, A4, A4i, B3, B4, B4i**)) and dorsolateral CSF-cNs (triangle, **Figure 3A3, A4, A4i, B3, B4, B4i**). In addition, we noticed *ptprna* expression in unknown spinal cells ventral to the central canal (asterisk, **Figure 3A3, A4, B1**).

**Figure 3.**
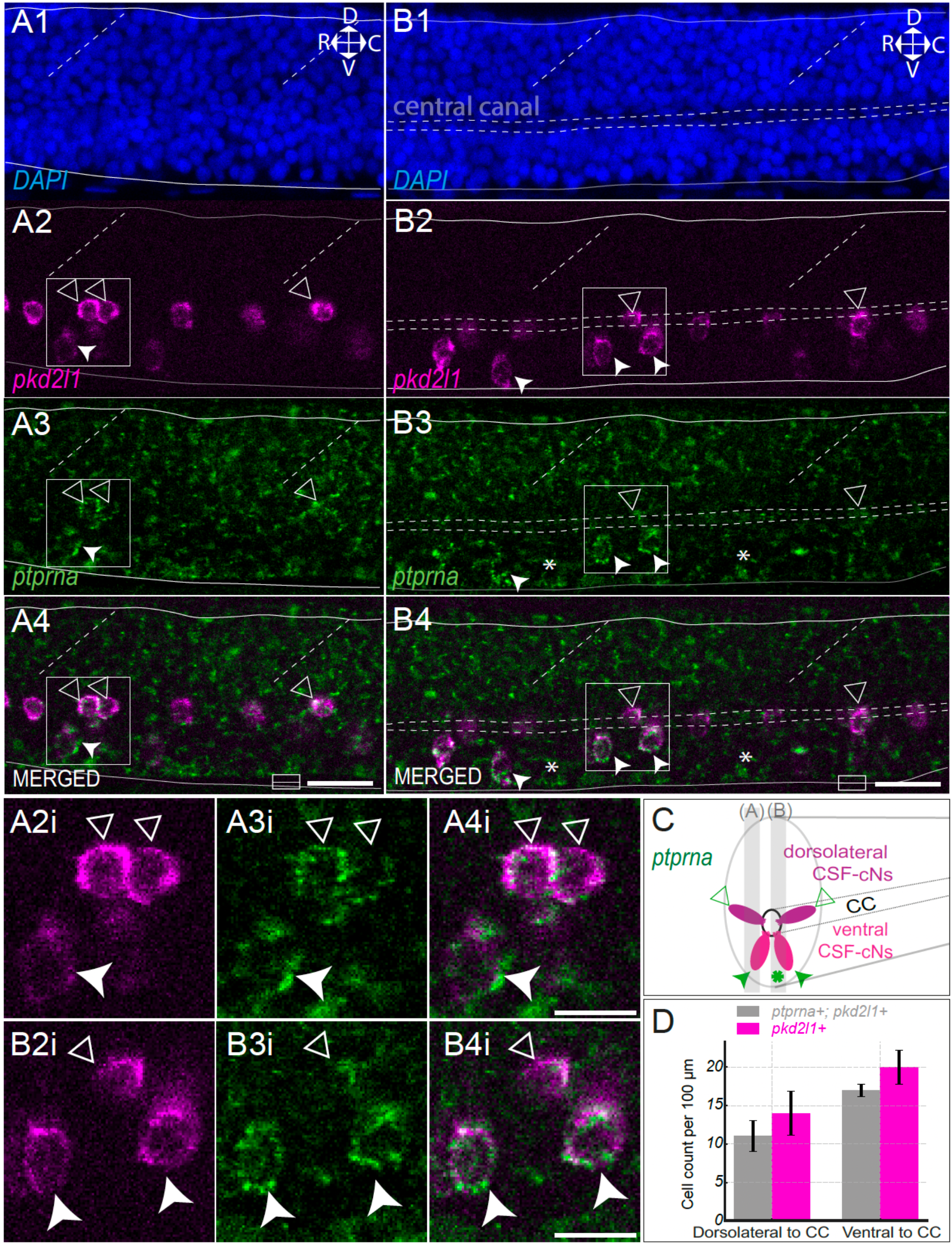
The receptor *ptprna* is expressed in ventral and dorsolateral spinal CSF-Ns. **(A)** Lateral sagittal optical section of the spinal cord of a 3 dpf larval zebrafish showing DAPI staining (A1) outlining dorsal and ventral boundaries (solid lines), the central canal (horizontal dashed lines), and somite boundaries (oblique dashed lines). Orientation: dorsal (D), ventral (V), rostral (R), caudal (C). *pkd2l1* labeling (A2, magenta) marks dorsolateral (triangle) and ventral (arrowhead) CSF-cNs. *ptprna* expression (A3, green) overlaps with *pkd2l1* in both dorsolateral and ventral CSF-cNs. Scale bar, 20 µm. **(A2i-A4i)** Magnified view of two dorsal (triangle) and one ventral (arrowhead) *pkd2l1*^+^/*ptprna*^+^ CSF-cNs. Scale bar, 10 µm. **(B)** Sagittal section showing DAPI staining (B1) outlining dorsal and ventral boundaries (solid lines), the central canal (horizontal dashed lines). *ptprna* expression (B3, green) is found in ventral (arrowhead) and dorsolateral (triangle) *pkd2l1*^+^ CSF-cNs (B2, magenta) and in *pkd2l1*^−^ cells near the central canal. Scale bar, 20 µm. **(B2i-B4i)** Magnified view of two ventral (arrowhead) and one dorsal *pkd2l1*^+^ *ptprna*^+^ CSF-cNs. Scale bar, 10 µm. **(C)** Schematic represents in 3D a transverse section of the spinal cord showing that both ventral and dorsolateral (magenta) CSF-cNs (*pkd2l1+)* express *ptprna* along with unknown cells around the central canal in the spinal cord (green asterisk). **(D)** Cell count per 100 µm of *ptprna*^+^ in CSF-cNs per larval (n = 3 larvae) are normalized per 100µm. In dorsolateral position to the central canal (CC): 10 ± 2.8 *ptprna*^+^/*pkd2l1*^+^ cells per 100 µm (grey); in ventral position to the central canal (CC): 15 ± 1.7 *ptprna*^+^/*pkd2l1*^+^ cells per 100 µm (grey). Cell count as *pkd2l1*^+^ cells (magenta): 14 ± 2.5 dorsolateral CSF-cNs per 100 µm and 19 ± 1.3 ventral CSF-cNs per 100 µm. Mean values are given ± s.e.m.

### Ldlrad2 is expressed in ventral and dorsolateral CSF-cNs and neighboring cells surrounding the central canal

To confirm the expression of *ldlrad2* in CSF-cNs, we performed HCR for *ldlrad2* and *pkd2l1* transcripts on 3 dpf AB *mifta*^-/-^ larvae (n = 6 fish, **Figure 4**). DAPI was added to delaminate the ventral and dorsal boundaries of the spinal cord and highlight the central canal (**Figure 4A1**). The *pkd2l1* receptor is expressed in both dorsolateral (triangle) and ventral (arrowhead) CSF-cNs (**Figure 4A2**). We observed all along the spinal cord co-expression of the receptor *ldlrad2* with *pkd2l1* (**Figure 4A3, A4**), indicating that dorsolateral (triangle, **Figure 4A2ii-A4ii**) and ventral (arrowhead) CSF-cNs (**Figure 4A2i-A4i**) express the *ldlrad2* receptor. In addition, we noticed that numerous other cells in contact with the cerebrospinal fluid (asterisk, **Figure 4A2i-A4i; 4A2ii-A4ii**) presumably ependymal radial glia cells (ERGs) (Jalalvand et al., 2014; Becker and Becker, 2022) also expressed the *ldlrad2* receptor (**Figure 4A3, A4, 4A2ii-A4ii**, asterisk).

**Figure 4.**
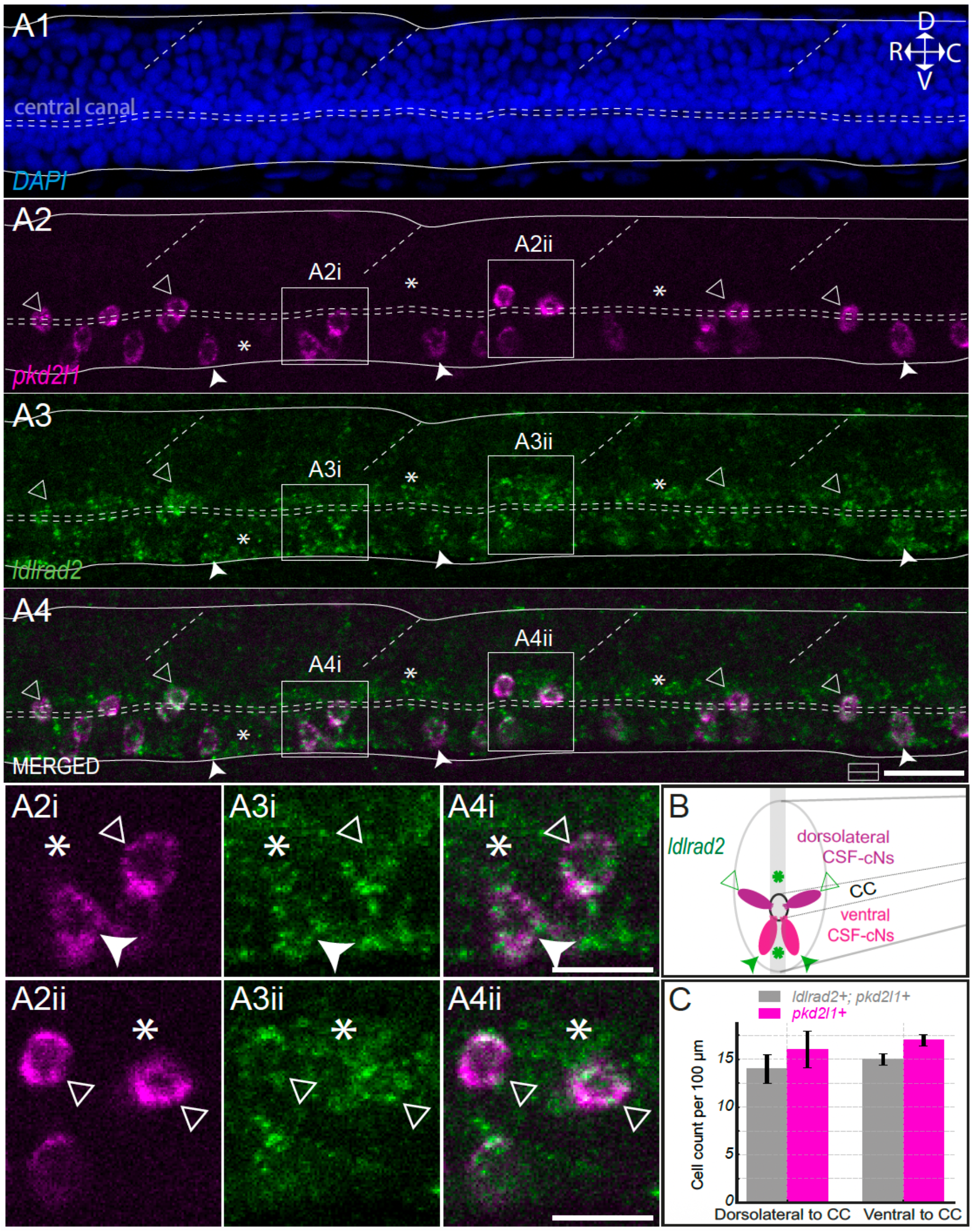
The receptor *ldlrad2* is expressed in ventral and dorsolateral CSF-cNs as well as other cells in contacting the cerebrospinal fluid, most likely corresponding to ependymal radial glia. **(A)** Sagittal optical section of the spinal cord showing DAPI staining (A1) outlining dorsal and ventral boundaries (solid lines), the central canal (horizontal dashed lines), and somite boundaries (oblique dashed lines). Orientation: dorsal (D), ventral (V), rostral (R), caudal (C). *pkd2l1* labeling (A2, magenta) marks dorsolateral (triangle) and ventral (arrowhead) CSF-cNs. *ldlrad2* expression (A3–A4, green) overlaps with *pkd2l1* in both dorsolateral and ventral CSF-cNs, and is also detected in other cells surrounding the central canal. Scale bar, 20 µm. **(A2i-A4i)** Magnified view of one dorsolateral and one ventral *pkd2l1*^+^/*ldlrad2*^+^ CSF-cNs and a *pkd2l1*^−^ in the ventral position to the central canal. **(A2ii-A4ii)** Magnified view of two dorsolateral *pkd2l1*^+^/*ldlrad2*^+^ CSF-cNs and *pkd2l1*^−^ cells surrounding the central canal (asterisk). (from box in A2–A4; scale bar, 10 µm). **(B)** Schematic represents in 3D a transverse section of the spinal cord showing that both ventral and dorsolateral (magenta) CSF-cNs (*pkd2l1+)* express *ldlrad2* along with unknown cells around the central canal and dorsally located in the spinal cord (green asterisk). **(C)** Cell counts of *ldlrad2+* per 100 µm in CSF-cNs per larval (n = 3 larvae) are normalized per 100µm. In dorsolateral position to the central canal (CC): 14 ± 1.5 *ldlrad2*^+^/*pkd2l1*^+^ cells per 100µm (grey); in ventral position to the central canal (CC): 15 ± 0.6 *ldlrad2*^+^/*pkd2l1*^+^ cells per 100µm (grey). Total *pkd2l1*^+^ cells (magenta): 16 ± 1.9 dorsolateral CSF-cNs per 100µm and 17 ± 0.6 ventral CSF-cNs per 100µm. Mean values are given ± s.e.m.

## DISCUSSION

In this study, we first confirmed expression of four chemoreceptors in CSF-cNs previously identified in the transcriptome from bulk population of CSF-cNs sorted by fluorescence (Prendergast et al., 2023): the metabotropic somatostatin receptor *sstr2a*, the metabotropic glutamate receptor *grm2a*, the receptor *ptprna* and the LDL receptor 2 *ldlrad2*.

### Receptor expression to specific cells contacting the CSF

While the *ldlrad2, grm2a* and *ptprna* receptors are present in all CSF-cNs, we find interesting specialization with specific receptors found in subtypes of CSF-cNs: the somatostatin receptor 2a is mainly expressed in ventral spinal CSF-cNs. This is particularly interesting as somatostatin 1 *sst1*.*1* is solely expressed by dorsal CSF-cNs: somatostatin1 could therefore coordinate the activity between dorsolateral and ventral CSF-cNs via the metabotropic somatostatin receptor 2a that acts as an inhibitor of secretion onto ventral CSF-cNs via the G protein Gαi (Rodrigues et al., 2018). The somatostatin receptor *sstr2a* was also expressed in dorsal spinal cord where it could modulate sensory processing such as itching in rodents (Flauaus et al., 2022) and chronic pain perception as shown previously via interaction with corticostatin (Morell et al., 2014) in addition to its role in psychiatric and neurodegenerative diseases (Beneyto et al., 2012; Ádori et al., 2015).

Furthermore, for most receptors investigated, sparse expression is also present in unknown spinal cells. For the LDL receptor 2 *ldlrad2*, the expression is also high in other cells surrounding the central canal that possibly correspond to ependymal radial glia (Becker and Becker 2014), which bear a motile cilium in the central canal (Bellegarda et al., 2023). Interestingly, the Reissner fiber in the central canal also exhibits multiple ligand-binding region of the LDL receptor family (Gobron et al., 1996; Sepúlveda et al., 2021).

Note that our list of receptors is of course not exhaustive. We selected some of the receptors that were only highly expressed and enriched in CSF-cNs versus all other cells of the trunk (Log Fold Change > 1.1). We could therefore have missed numerous receptors not enriched in CSF-cNs, highly expressed in other cells of the trunk (such as muscles, notochord, gut), or just expressed at low levels in CSF-cNs.

### Physiological context: when does the CSF change composition and could modulate these receptors?

In fish, lamprey and mouse, CSF-cNs have been involved in diverse physiological functions in the context of motor control, posture and locomotion (Wyart et al., 2009; Hubbard et al., 2016; Bohm et al., 2016; Quan et al., 2020; Wu et al., 2021; Gerstmann et al., 2022; Nakamura et al., 2023; Jalalvand et al., 2016a; Jalalvand et al., 2016b). In addition, this interoceptive system has been also involved in morphogenesis to strengthen the body axis and spine (Sternberg et al., 2018; Cantaut-Belarif et al., 2018; Zhang et al., 2018; Bearce et al., 2022; Gaillard et al., 2023), detection of pathogen invasion to enhance host defence (Prendergast et al., 2023) as well as modulation of the stem cells niche after spinal cord injury (Wyart et al., 2023; Kathe et al., 2022; Yue et al. 2024).

Our data here suggest that somatostatin may act as a signaling molecule from dorsolateral to ventral CSF-cNs. Dorsolateral CSF-cNs specifically express somatostatin (Djenoune et al., 2017). During locomotion, the tail bends from side to side, which activates ipsilateral dorsolateral CSF-cNs that respond to left / right curvature of the trunk (Böhm et al., 2016), which can dampen the oscillations and reduce bout duration (Quan et al., 2020). We show here that ventral CSF-cNs express the somatostatin receptors *sstr2a*. Therefore, upon activation of dorsolateral CSF-cNs during locomotion, somatostatin could act to reduce intracellular calcium in ventral CSF-cNs, and thereby decreasing the release of numerous other peptides and secreted proteins such as *urp1, urp2, msmp2*, or *nppc* (Quan et al., 2015; Prendergast et al., 2023).

CSF lipoproteins transport lipids and associated proteins that contribute to lipid homeostasis in the central nervous system and support neural development (Tsujita et al., 2024; Merrill et al. 2023). This pathway could be relevant to morphogenesis regulated by CSF-cNs and the Reissner fiber (Wyart et al., 2023). The Reissner fiber is indeed formed by the glycoprotein SCO-spondin, which contains low-density lipoprotein receptor type A (LDLrA) domains (Meiniel and Meiniel, 2007). *In vitro* experiments have shown that LDL can bind SCO-spondin and modulate its neurogenic activity (Vera et al., 2015). In addition, LDL-family particles can carry signaling molecules, including morphogens from the Hedgehog (Hh) and Wnt families (Panáková et al., 2005; Willnow et al., 2007). SCO-spondin has been proposed to facilitate morphogen distribution in the CSF by interacting with lipoproteins (Vera et al., 2015). LDL receptor 2a (*ldlr2a*) appears to be expressed by cells in contact with the cerebrospinal fluid, including CSF-cNs, and most likely ependymal radial glia and floor plate cells. These cells could therefore use LDLR2a to capture LDL/associated cargo (lipids and/or morphogens) from the CSF, potentially facilitated by interactions with the Reissner fiber during morphogenesis.

During spinal cord injury or when pathogens invade the CSF during an infection of the central nervous system, glutamate can be released from the cytoplasm following cell death. In human bacterial meningitis, elevated CSF glutamate levels have been associated with clinical outcomes (Spranger et al., 1996). One potential pathway is that excess glutamate is detected by CSF-cNs via activation of this Gi/o-coupled grm2a receptor. Receptor activation inhibits adenylyl cyclase (AC), reduces intracellular cAMP, and can suppress voltage-gated calcium channel (VGCC) activity, thereby decreasing Ca^2+^ influx (Reiner and Levitz, 2018). Such a reduction in Ca^2+^ entry could be neuroprotective by limiting Ca^2+^-dependent cellular damage under inflammatory conditions. Consistent with reduced excitability, intraventricular injection of *Streptococcus pneumoniae* in zebrafish silenced spontaneous global Ca^2+^ activity in CSF-cNs (Prendergast et al., 2023). In the context of spinal cord injury, glutamate may be sensed by CSF-cNs via the metabotropic glutamate receptor Grm2a, which can promote GABA release (Corti et al., 2007) and modulate the stem cell niche (New et al., 2023). Because infection and injury converge on excitotoxic and inflammatory cascades, Grm2a-dependent CSF-cN signaling may couple acute changes in excitability to longer-term repair programs, consistent with regenerative responses reported in mice (Kathe et al., 2022; Yue et al., 2024).

Ptprn is localized to dense-core secretory vesicles in neuroendocrine cells and is involved in secretory granule biogenesis and regulation of secretion (Cai et al., 2011). CSF-cNs are secretory cells containing dense-core vesicles (Djenoune et al., 2017) and express numerous genes encoding neuropeptides and other secreted factors implicated in locomotion, morphogenesis and innate immunity (Prendergast et al., 2023; Wyart et al., 2023). The expression of *ptprna* in CSF-cNs therefore suggests that this receptor contributes to the regulation of secretory machinery and may participate in controlling the release of bioactive compounds into the cerebrospinal fluid.

Altogether, our study opens new paths for investigation of chemosensory signaling in the spinal cord and at the interface with the cerebrospinal fluid that can carry long range signaling between the nervous system and other systems and organs in the body.

## Materials and Methods

### Zebrafish housing, mutant and strain

Animal handling and procedures were validated by the Paris Brain Institute (ICM) and the French National Ethics Committee (Comité National de Réflexion Éthique sur l’Expérimentation Animale; APAFIS no. 2018071217081175 and APAFIS no. #38209-2022080517223947 and APAFIS no. #51743-2024102515029150) in agreement with European Union legislation. To avoid pigmentation, all experiments were performed on *Danio rerio* larvae of AB strain carrying the *mitfa*^−*/*−^ mutation. Adult zebrafish were reared at a maximum density of six animals per liter in a 14/10 h light–dark cycle environment at 28.5 °C. Larval zebrafish were typically raised in Petri dishes filled with system water under the same conditions in terms of temperature and lighting as for adults.

### Fluorescent *in situ* hybridization using high chain reaction (HCR)

HCR probes were custom-designed and synthesized by Molecular Instruments (Los Angeles, CA, USA) based on NCBI mRNA reference sequences: *pkd2l1* (XM_690312), *sstr2a* (XM_005170121.5), *ldlrad2* (XM_003199510.6), *ptprna* (XM_017357652.3), and *grm2a* boosted version (XM_005166117.4). HCR reagents, including hairpins and buffers, were obtained from the same supplier. Double-staining HCR was performed for all experiments using a probe for *pkd2l1* and a probe for the receptor of interest. The HCR hairpins were labeled with Alexa Fluor 488, Alexa Fluor 546, or Alexa Fluor 647. The protocol of « whole-mount zebrafish embryos and larvae » used for HCR was modified based on Shainer et al., Science Advances 2022. Briefly, embryos/larvae (1–10 per mL) were fixed in 1 mL freshly prepared 4% paraformaldehyde (PFA) in Dulbecco’s phosphate-buffered saline (DPBS) overnight at 4 °C with gentle shaking to preserve tissue integrity. Fixed samples were washed 3 × 5 min in DPBST (DPBS + 0.1% Tween 20) and permeabilized for 10 min in prechilled 100% methanol at −20 °C. For rehydration, embryos/larvae were washed for 5 min in 50% methanol/50% DPBST, 5 min in 25% methanol/75% DPBST, and 5 × 5 min in DPBST. To optimize the probe permeabilization, 2-dpf embryos were treated with 10 µg/mL Proteinase K in 1× DPBST for 20 min and 3-dpf larvae for 30 min at room temperature, followed by post-fixation in 4% PFA for 20 min. To increase the signaling for weakly expressed metabotropic receptor genes, the concentration of the probe was prepared for 32 or 64nM in final solutions (8 or 16 μl of 1 μM probe stock). The standard concentration 16nM was used for *pkd2l1* (4 μl of 1 μM probe stock). After prehybridization in 250 μL prewarmed hybridization buffer for 30 min at 37 °C, embryos/larvae in probe solution were incubated 12–16 h at 37 °C. Excess probes were removed by four 30 min washes in 250 μL prewarmed probe wash buffer at 37 °C, followed by two 20 min washes in 5× SSCT (sodium chloride sodium citrate + 0.1% Tween 20) at room temperature. Embryos/larvae were preamplified in 250 μL amplification buffer for 30 min at room temperature. Hairpins h1 and h2 (60 pmol each) were prepared by snap-cooling 10 μL of 3 μM stock at 95 °C for 90 s, then cooling in the dark for 30 min. For double labeling, 5 μL of each hairpin and corresponding amplifiers were included. Hairpins were combined in a 250 μL amplification buffer with DAPI, and embryos/larvae were incubated in this solution overnight (12– 16 h) in the dark at room temperature. Following removal of the amplification buffer, embryos/larvae were incubated in hairpin solution containing DAPI (1:1000) for 12–16 h in the dark at room temperature. The next day, excess hairpins were removed by 3 × 20 min washes in 500 μL 5× SSCT at room temperature. Samples were mounted in ibidi Mounting Medium (Ref: 50001, ibidi GmbH, Germany) for imaging.

### Confocal Imaging

Imaging was performed on a Leica SP8 DLS (inverted) or SP8 X White Light Laser confocal microscope (Leica Microsystems, Wetzlar, Germany) equipped with a 40 X oil-immersion objective (NA = 1.3). Laser lines used were 405, 488, 552, and 638 nm (SP8 DLS) or 405 nm and a white laser (470–670 nm; SP8 X), applied in sequential scanning mode to avoid spectral bleed-through. Emission windows were 415–465 nm (DAPI), 498–520 nm (488 nm excitation), 568– 600 nm (546 nm excitation), and ≥657 nm (647 nm excitation). Image acquisition was performed with LAS X software (Leica Microsystems) and processed in Fiji (https://imagej.net/software/fiji/downloads, Schindelin et al., 2012).

### Image Quantification

We quantified all cells expressing the receptors and identified whether they were ventral or dorsolateral CSF-cNs, other CSF-contacting cells (floor plate, roof plate, ependymal radial glia) or other unknown cells in the dorsal spinal cord. Three segments, each comprising four somites in the rostral, middle, and caudal regions of the trunk, were imaged within a single larva. For each segment, Z-stacks were acquired from 30–40 μm-thick sections using a step size of 1 μm, resulting in 30–40 slices with a field of view measuring 291 × 145 μm. The *pkd2l1*+ cells were distinguished in two groups: *(i)* Ventral CSF-cNs: a ventral row of cells adjacent to the central canal *(ii)* dorsolateral CSF-cNs: a dorsal row of cells adjacent to the central canal. The receptor-expressing cells were counted in ventral and dorsolateral CSF-cNs and in other *pkd2l1*-cells. The proportion of receptor-expressing cells among *pkd2l1*+ cells was assessed separately for ventral and dorsolateral CSF-cNs. The proportion of receptor-expressing cells among other cell types was calculated separately for the dorsal-to-central canal region and the P3 domain. Cells were counted per 100 µm for each location along the rostrocaudal axis (rostral, middle, and caudal) within a single larva. For each receptor, three double-labeled larvae were analyzed.

### Statistics

All values provided in the text are given as mean +/- standard deviation.

## Acknowledgments

We would like to thank for their inputs and camaraderie spirit the Wyart lab “SIBBIL” (https://wyartlab.org; https://parisbraininstitute.org/paris-brain-institute-research-teams/sibbil-navigation-sensorimotor-integration-brain-body-integration-lab) for providing great insight along the way, particularly Dr. Giulia Messa, Dr. Emma Partiot, Dr. Clothilde Colart for constructive discussions that helped shape the manuscript. We thank A. Arneau, N. Jezequel, C. Lejeune, S. Nunes Figueiredo, B. Daboval of the core facility Pheno-Zfish for fish care. We thank the 2020 Fondation Bettencourt-Schueller (FBS-don-0031) award (“Identity and organization or neuronal networks controlling exploration”), a New York Stem Cell Foundation (NYSCF) Robertson Award 2016 research grant (NYSCF-R-NI39), the 2020 Prize Equipe ‘Fondation pour la Recherche Médicale’ (FRM-EQU202003010612) ‘Neuronal circuits underlying navigation: from genes to behavioral models’, a 2020 European Research Council Consolidator grant no. 101002870 (2021– 2026) ‘Exploratome: Circuit mechanisms underlying sensory-evoked navigation’ and a National Institutes of Health grant no. 1U19NS104653-01 awarded to C.W., the European Union’s Horizon 2020 research and innovation programme under a Marie Skłodowska-Curie grant no. 813457 awarded to C.W. The project benefited as well from the support of the Agence Nationale pour la Recherche (ANR) ANR-22-CE37-0023 named LOCOCONNECT, ANR-23-CE16-0017-02 named RocSMAP, ANR-24-CE16-7992 named CIRCOLOCO, ANR-21-CE14-0042 named MOTOMYO and ANR-21-CE13-0008 named ASCENTS.

## Author Contributions

Conceptualization: CW & FQ; HCR execution and optimization: EV, LM, LT; Quantification: EV; Analysis: EV; Data visualization: EV; Writing of initial draft: CW, FQ, EV; Supervision: CW & FQ; Funding: CW.

## Classification

Biological Sciences, Neurosciences

## References

Ádori, C., Glück, L., Barde, S., Yoshitake, T., Kovacs, G. G., Mulder, J., Maglóczky, Z., Havas, L., Bölcskei, K., Mitsios, N., Uhlén, M., Szolcsányi, J., Kehr, J., Rönnbäck, A., Schwartz, T., Rehfeld, J. F., Harkany, T., Palkovits, M., Schulz, S., & Hökfelt, T. (2015). Critical role of somatostatin receptor 2 in the vulnerability of the central noradrenergic system: New aspects on Alzheimer’s disease. Acta Neuropathologica, 129(4), 541–563. 10.1007/s00401-015-1394-3

Agduhr, E. (1922). Über ein zentrales Sinnesorgan (?) bei den Vertebraten. Zeitschrift für Anatomie und Entwicklungsgeschichte, 66(3–6), 223–360. 10.1007/BF02593586

Bearce, E. A., Irons, Z. H., O’Hara-Smith, J. R., Kuhns, C. J., Fisher, S. I., Crow, W. E., & Grimes, D. T. (2022). Urotensin II-related peptides, Urp1 and Urp2, control zebrafish spine morphology. eLife, 11, e83883. 10.7554/eLife.83883

Becker, T., & Becker, C. G. (2014). Axonal regeneration in zebrafish. Current Opinion in Neurobiology, 27, 186–191. 10.1016/j.conb.2014.03.019

Becker, T., & Becker, C. G. (2022). Regenerative neurogenesis: The integration of developmental, physiological and immune signals. Development (Cambridge, England), 149(8), dev199907. 10.1242/dev.199907

Bellegarda, C., Zavard, G., Moisan, L., Brochard-Wyart, F., Joanny, J.-F., Gray, R. S., Cantaut-Belarif, Y., & Wyart, C. (2023). The Reissner fiber under tension in vivo shows dynamic interaction with ciliated cells contacting the cerebrospinal fluid. eLife, 12, e86175. 10.7554/eLife.86175

Beneyto, M., Morris, H. M., Rovensky, K. C., & Lewis, D. A. (2012). Lamina- and cell-specific alterations in cortical somatostatin receptor 2 mRNA expression in schizophrenia. Neuropharmacology, 62(3), 1598–1605. 10.1016/j.neuropharm.2010.12.029

Böhm, U. L., Prendergast, A., Djenoune, L., Nunes Figueiredo, S., Gomez, J., Stokes, C., Kaiser, S., Suster, M., Kawakami, K., Charpentier, M., Concordet, J.-P., Rio, J.-P., Del Bene, F., & Wyart, C. (2016). CSF-contacting neurons regulate locomotion by relaying mechanical stimuli to spinal circuits. Nature Communications, 7, 10866. 10.1038/ncomms10866

Cai, T., Hirai, H., Zhang, G., Zhang, M., Takahashi, N., Kasai, H., Satin, L. S., Leapman, R. D., & Notkins, A. L. (2011). Deletion of Ia-2 and/or Ia-2β in mice decreases insulin secretion by reducing the number of dense core vesicles. Diabetologia, 54(9), 2347–2357. 10.1007/s00125-011-2221-6

Cantaut-Belarif, Y., Sternberg, J. R., Thouvenin, O., Wyart, C., & Bardet, P.-L. (2018). The Reissner Fiber in the Cerebrospinal Fluid Controls Morphogenesis of the Body Axis. Current Biology: CB, 28(15), 2479–2486.e4. 10.1016/j.cub.2018.05.079

Corti, C., Battaglia, G., Molinaro, G., Riozzi, B., Pittaluga, A., Corsi, M., Mugnaini, M., Nicoletti, F., & Bruno, V. (2007). The use of knock-out mice unravels distinct roles for mGlu2 and mGlu3 metabotropic glutamate receptors in mechanisms of neurodegeneration/neuroprotection. The Journal of Neuroscience: The Official Journal of the Society for Neuroscience, 27(31), 8297–8308. 10.1523/JNEUROSCI.1889-07.2007

Dale, N., Roberts, A., Ottersen, O. P., & Storm-Mathisen, J. (1987). The morphology and distribution of “Kolmer-Agduhr cells”, a class of cerebrospinal-fluid-contacting neurons revealed in the frog embryo spinal cord by GABA immunocytochemistry. Proceedings of the Royal Society of London. Series B, Biological Sciences, 232(1267), 193–203. 10.1098/rspb.1987.0068

Di Bella, D. J., Carcagno, A. L., Bartolomeu, M. L., Pardi, M. B., Löhr, H., Siegel, N., Hammerschmidt, M., Marín-Burgin, A., & Lanuza, G. M. (2019). Ascl1 Balances Neuronal versus Ependymal Fate in the Spinal Cord Central Canal. Cell Reports, 28(9), 2264–2274.e3. 10.1016/j.celrep.2019.07.087

Djenoune, L., Desban, L., Gomez, J., Sternberg, J. R., Prendergast, A., Langui, D., Quan, F. B., Marnas, H., Auer, T. O., Rio, J.-P., Del Bene, F., Bardet, P.-L., & Wyart, C. (2017). The dual developmental origin of spinal cerebrospinal fluid-contacting neurons gives rise to distinct functional subtypes. Scientific Reports, 7(1), 719. 10.1038/s41598-017-00350-1

Djenoune, L., Khabou, H., Joubert, F., Quan, F. B., Nunes Figueiredo, S., Bodineau, L., Del Bene, F., Burcklé, C., Tostivint, H., & Wyart, C. (2014). Investigation of spinal cerebrospinal fluid-contacting neurons expressing PKD2L1: Evidence for a conserved system from fish to primates. Frontiers in Neuroanatomy, 8, 26. 10.3389/fnana.2014.00026

Djenoune, L., & Wyart, C. (2017). Light on a sensory interface linking the cerebrospinal fluid to motor circuits in vertebrates. Journal of Neurogenetics, 31(3), 113–127. 10.1080/01677063.2017.1359833

Fagan, A. M., Henson, R. L., Li, Y., Boerwinkle, A. H., Xiong, C., Bateman, R. J., Goate, A., Ances, B. M., Doran, E., Christian, B. T., Lai, F., Rosas, H. D., Schupf, N., Krinsky-McHale, S., Silverman, W., Lee, J. H., Klunk, W. E., Handen, B. L., Allegri, R. F., … Dominantly Inherited Alzheimer Network. (2021). Comparison of CSF biomarkers in Down syndrome and autosomal dominant Alzheimer’s disease: A cross-sectional study. The Lancet. Neurology, 20(8), 615–626. 10.1016/S1474-4422(21)00139-3

Fidelin, K., Djenoune, L., Stokes, C., Prendergast, A., Gomez, J., Baradel, A., Del Bene, F., & Wyart, C. (2015). State-Dependent Modulation of Locomotion by GABAergic Spinal Sensory Neurons. Current Biology: CB, 25(23), 3035–3047. 10.1016/j.cub.2015.09.070

Flauaus, C., Engel, P., Zhou, F., Petersen, J., Ruth, P., Lukowski, R., Schmidtko, A., & Lu, R. (2022). Slick Potassium Channels Control Pain and Itch in Distinct Populations of Sensory and Spinal Neurons in Mice. Anesthesiology, 136(5), 802–822. 10.1097/ALN.0000000000004163

Gaillard, A.-L., Mohamad, T., Quan, F. B., de Cian, A., Mosimann, C., Tostivint, H., & Pézeron, G. (2023). Urp1 and Urp2 act redundantly to maintain spine shape in zebrafish larvae. Developmental Biology, 496, 36–51. 10.1016/j.ydbio.2023.01.010

Gerstmann, K., Jurčić, N., Blasco, E., Kunz, S., de Almeida Sassi, F., Wanaverbecq, N., & Zampieri, N. (2022). The role of intraspinal sensory neurons in the control of quadrupedal locomotion. Current Biology: CB, 32(11), 2442–2453.e4. 10.1016/j.cub.2022.04.019

Gobron, S., Monnerie, H., Meiniel, R., Creveaux, I., Lehmann, W., Lamalle, D., Dastugue, B., & Meiniel, A. (1996). SCO-spondin: A new member of the thrombospondin family secreted by the subcommissural organ is a candidate in the modulation of neuronal aggregation. Journal of Cell Science, 109 (Pt 5), 1053–1061. 10.1242/jcs.109.5.1053

Hermann, P., Appleby, B., Brandel, J.-P., Caughey, B., Collins, S., Geschwind, M. D., Green, A., Haïk, S., Kovacs, G. G., Ladogana, A., Llorens, F., Mead, S., Nishida, N., Pal, S., Parchi, P., Pocchiari, M., Satoh, K., Zanusso, G., & Zerr, I. (2021). Biomarkers and diagnostic guidelines for sporadic Creutzfeldt-Jakob disease. The Lancet. Neurology, 20(3), 235–246. 10.1016/S1474-4422(20)30477-4

Hökfelt, T., Fuxe, K., & Goldstein, M. (1973). Immunohistochemical localization of aromatic L-amino acid decarboxylase (DOPA decarboxylase) in central dopamine and 5-hydroxytryptamine nerve cell bodies of the rat. Brain Research, 53(1), 175–180. 10.1016/0006-8993(73)90776-2

Huang, A. L., Chen, X., Hoon, M. A., Chandrashekar, J., Guo, W., Tränkner, D., Ryba, N. J. P., & Zuker, C. S. (2006). The cells and logic for mammalian sour taste detection. Nature, 442(7105), 934–938. 10.1038/nature05084

Huang, P., Xiong, F., Megason, S. G., & Schier, A. F. (2012). Attenuation of Notch and Hedgehog signaling is required for fate specification in the spinal cord. PLoS Genetics, 8(6), e1002762. 10.1371/journal.pgen.1002762

Hubbard, J. M., Böhm, U. L., Prendergast, A., Tseng, P.-E. B., Newman, M., Stokes, C., & Wyart, C. (2016). Intraspinal Sensory Neurons Provide Powerful Inhibition to Motor Circuits Ensuring Postural Control during Locomotion. Current Biology: CB, 26(21), 2841–2853. 10.1016/j.cub.2016.08.026

Jaeger, C. B., Teitelman, G., Joh, T. H., Albert, V. R., Park, D. H., & Reis, D. J. (1983). Some neurons of the rat central nervous system contain aromatic-L-amino-acid decarboxylase but not monoamines. Science (New York, N.Y.), 219(4589), 1233–1235. 10.1126/science.6131537

Jalalvand, E., Robertson, B., Tostivint, H., Wallén, P., & Grillner, S. (2016a). The Spinal Cord Has an Intrinsic System for the Control of pH. Current Biology: CB, 26(10), 1346–1351. 10.1016/j.cub.2016.03.048

Jalalvand, E., Robertson, B., Wallén, P., & Grillner, S. (2016b). Ciliated neurons lining the central canal sense both fluid movement and pH through ASIC3. Nature Communications, 7, 10002. 10.1038/ncomms10002

Jalalvand, E., Robertson, B., Wallén, P., Hill, R. H., & Grillner, S. (2014). Laterally projecting cerebrospinal fluid-contacting cells in the lamprey spinal cord are of two distinct types. The Journal of Comparative Neurology, 522(8), 1753–1768. 10.1002/cne.23542

Kathe, C., Skinnider, M. A., Hutson, T. H., Regazzi, N., Gautier, M., Demesmaeker, R., Komi, S., Ceto, S., James, N. D., Cho, N., Baud, L., Galan, K., Matson, K. J. E., Rowald, A., Kim, K., Wang, R., Minassian, K., Prior, J. O., Asboth, L., … Courtine, G. (2022). The neurons that restore walking after paralysis. Nature, 611(7936), 540–547. 10.1038/s41586-022-05385-7

Kolmer, W. (1921). Das „Sagittalorgan” der Wirbeltiere. Zeitschrift für Anatomie und Entwicklungsgeschichte, 60(3–6), 652–717. 10.1007/BF02593657

Marie-Hardy, L., Slimani, L., Messa, G., El Bourakkadi, Z., Prigent, A., Sayetta, C., Koëth, F., Pascal-Moussellard, H., Wyart, C., & Cantaut-Belarif, Y. (2023). Loss of CSF-contacting neuron sensory function is associated with a hyper-kyphosis of the spine reminiscent of Scheuermann’s disease. Scientific Reports, 13(1), 5529. 10.1038/s41598-023-32536-1

Meiniel, O., & Meiniel, A. (2007). The complex multidomain organization of SCO-spondin protein is highly conserved in mammals. Brain Research Reviews, 53(2), 321–327. 10.1016/j.brainresrev.2006.09.007

Merrill, N. J., Davidson, W. S., He, Y., Díaz Ludovico, I., Sarkar, S., Berger, M. R., McDermott, J. E., Van Eldik, L. J., Wilcock, D. M., Monroe, M. E., Kyle, J. E., Bruce, K. D., Heinecke, J. W., Vaisar, T., Raber, J., Quinn, J. F., & Melchior, J. T. (2023). Human cerebrospinal fluid contains diverse lipoprotein subspecies enriched in proteins implicated in central nervous system health. Science Advances, 9(35), eadi5571. 10.1126/sciadv.adi5571

Morell, M., Camprubí-Robles, M., Culler, M. D., de Lecea, L., & Delgado, M. (2014). Cortistatin attenuates inflammatory pain via spinal and peripheral actions. Neurobiology of Disease, 63, 141–154. 10.1016/j.nbd.2013.11.022

Nakamura, Y., Kurabe, M., Matsumoto, M., Sato, T., Miyashita, S., Hoshina, K., Kamiya, Y., Tainaka, K., Matsuzawa, H., Ohno, N., & Ueno, M. (2023). Cerebrospinal fluid-contacting neuron tracing reveals structural and functional connectivity for locomotion in the mouse spinal cord. eLife, 12, e83108. 10.7554/eLife.83108

New, L. E., Yanagawa, Y., McConkey, G. A., Deuchars, J., & Deuchars, S. A. (2023). GABAergic regulation of cell proliferation within the adult mouse spinal cord. Neuropharmacology, 223, 109326. 10.1016/j.neuropharm.2022.109326

Orts-Del’Immagine, A., Cantaut-Belarif, Y., Thouvenin, O., Roussel, J., Baskaran, A., Langui, D., Koëth, F., Bivas, P., Lejeune, F.-X., Bardet, P.-L., & Wyart, C. (2020). Sensory Neurons Contacting the Cerebrospinal Fluid Require the Reissner Fiber to Detect Spinal Curvature In Vivo. Current Biology: CB, 30(5), 827–839.e4. 10.1016/j.cub.2019.12.071

Orts-Del’Immagine, A., Wanaverbecq, N., Tardivel, C., Tillement, V., Dallaporta, M., & Trouslard, J. (2012). Properties of subependymal cerebrospinal fluid contacting neurones in the dorsal vagal complex of the mouse brainstem. The Journal of Physiology, 590(16), 3719–3741. 10.1113/jphysiol.2012.227959

Panáková, D., Sprong, H., Marois, E., Thiele, C., & Eaton, S. (2005). Lipoprotein particles are required for Hedgehog and Wingless signalling. Nature, 435(7038), 58–65. 10.1038/nature03504

Parnetti, L., Gaetani, L., Eusebi, P., Paciotti, S., Hansson, O., El-Agnaf, O., Mollenhauer, B., Blennow, K., & Calabresi, P. (2019). CSF and blood biomarkers for Parkinson’s disease. The Lancet. Neurology, 18(6), 573–586. 10.1016/S1474-4422(19)30024-9

Petracca, Y. L., Sartoretti, M. M., Di Bella, D. J., Marin-Burgin, A., Carcagno, A. L., Schinder, A. F., & Lanuza, G. M. (2016). The late and dual origin of cerebrospinal fluid-contacting neurons in the mouse spinal cord. Development, 143(5), 880–891. 10.1242/dev.129254

Prendergast, A. E., Jim, K. K., Marnas, H., Desban, L., Quan, F. B., Djenoune, L., Laghi, V., Hocquemiller, A., Lunsford, E. T., Roussel, J., Keiser, L., Lejeune, F.-X., Dhanasekar, M., Bardet, P.-L., Levraud, J.-P., van de Beek, D., Vandenbroucke-Grauls, C. M. J. E., & Wyart, C. (2023). CSF-contacting neurons respond to Streptococcus pneumoniae and promote host survival during central nervous system infection. Current Biology: CB, 33(5), 940–956.e10. 10.1016/j.cub.2023.01.039

Quan, F. B., Desban, L., Mirat, O., Kermarquer, M., Roussel, J., Koëth, F., Marnas, H., Djenoune, L., Lejeune, F.-X., Tostivint, H., & Wyart, C. (2020). Somatostatin 1.1 contributes to the innate exploration of zebrafish larva. Scientific Reports, 10(1), 15235. 10.1038/s41598-020-72039-x

Quan, F. B., Dubessy, C., Galant, S., Kenigfest, N. B., Djenoune, L., Leprince, J., Wyart, C., Lihrmann, I., & Tostivint, H. (2015). Comparative distribution and in vitro activities of the urotensin II-related peptides URP1 and URP2 in zebrafish: Evidence for their colocalization in spinal cerebrospinal fluid-contacting neurons. PloS One, 10(3), e0119290. 10.1371/journal.pone.0119290

Reiner, A., & Levitz, J. (2018). Glutamatergic Signaling in the Central Nervous System: Ionotropic and Metabotropic Receptors in Concert. Neuron, 98(6), 1080–1098. 10.1016/j.neuron.2018.05.018

Reissner, E. (1860). Beiträge zur Kenntniss vom Bau des Rückenmarkes von Petromyzon fluviatilis L. 545–588.

Ren, L.-Q., Chen, M., Hultborn, H., Guo, S., Zhang, Y., & Zhang, M. (2017). Heterogenic Distribution of Aromatic L-Amino Acid Decarboxylase Neurons in the Rat Spinal Cord. Frontiers in Integrative Neuroscience, 11, 31. 10.3389/fnint.2017.00031

Rodriguez, M., Frost, J. A., & Schonbrunn, A. (2018). Real-Time Signaling Assays Demonstrate Somatostatin Agonist Bias for Ion Channel Regulation in Somatotroph Tumor Cells. Journal of the Endocrine Society, 2(7), 779–793. 10.1210/js.2018-00115

Rose, C. D., Pompili, D., Henke, K., Van Gennip, J. L. M., Meyer-Miner, A., Rana, R., Gobron, S., Harris, M. P., Nitz, M., & Ciruna, B. (2020). SCO-Spondin Defects and Neuroinflammation Are Conserved Mechanisms Driving Spinal Deformity across Genetic Models of Idiopathic Scoliosis. Current Biology: CB, 30(12), 2363–2373.e6. 10.1016/j.cub.2020.04.020

Schindelin, J., Arganda-Carreras, I., Frise, E., Kaynig, V., Longair, M., Pietzsch, T., Preibisch, S., Rueden, C., Saalfeld, S., Schmid, B., Tinevez, J.-Y., White, D. J., Hartenstein, V., Eliceiri, K., Tomancak, P., & Cardona, A. (2012). Fiji: An open-source platform for biological-image analysis. Nature Methods, 9(7), 676–682. 10.1038/nmeth.2019

Sepúlveda, V., Maurelia, F., González, M., Aguayo, J., & Caprile, T. (2021). SCO-spondin, a giant matricellular protein that regulates cerebrospinal fluid activity. Fluids and Barriers of the CNS, 18(1), 45. 10.1186/s12987-021-00277-w

Shin, J., Poling, J., Park, H.-C., & Appel, B. (2007). Notch signaling regulates neural precursor allocation and binary neuronal fate decisions in zebrafish. Development (Cambridge, England), 134(10), 1911–1920. 10.1242/dev.001602

Spranger, M., Schwab, S., Krempien, S., Winterholler, M., Steiner, T., & Hacke, W. (1996). Excess glutamate levels in the cerebrospinal fluid predict clinical outcome of bacterial meningitis. Archives of Neurology, 53(10), 992–996. 10.1001/archneur.1996.00550100066016

Sternberg, J. R., Prendergast, A. E., Brosse, L., Cantaut-Belarif, Y., Thouvenin, O., Orts-Del’Immagine, A., Castillo, L., Djenoune, L., Kurisu, S., McDearmid, J. R., Bardet, P.-L., Boccara, C., Okamoto, H., Delmas, P., & Wyart, C. (2018). Pkd2l1 is required for mechanoception in cerebrospinal fluid-contacting neurons and maintenance of spine curvature. Nature Communications, 9(1), 3804. 10.1038/s41467-018-06225-x

Stoeckel, M.-E., Uhl-Bronner, S., Hugel, S., Veinante, P., Klein, M.-J., Mutterer, J., Freund-Mercier, M.-J., & Schlichter, R. (2003). Cerebrospinal fluid-contacting neurons in the rat spinal cord, a gamma-aminobutyric acidergic system expressing the P2X2 subunit of purinergic receptors, PSA-NCAM, and GAP-43 immunoreactivities: Light and electron microscopic study. The Journal of Comparative Neurology, 457(2), 159–174. 10.1002/cne.10565

Thouvenin, O., Keiser, L., Cantaut-Belarif, Y., Carbo-Tano, M., Verweij, F., Jurisch-Yaksi, N., Bardet, P.-L., van Niel, G., Gallaire, F., & Wyart, C. (2020). Origin and role of the cerebrospinal fluid bidirectional flow in the central canal. eLife, 9, e47699. 10.7554/eLife.47699

Troutwine, B. R., Gontarz, P., Konjikusic, M. J., Minowa, R., Monstad-Rios, A., Sepich, D. S., Kwon, R. Y., Solnica-Krezel, L., & Gray, R. S. (2020). The Reissner Fiber Is Highly Dynamic In Vivo and Controls Morphogenesis of the Spine. Current Biology: CB, 30(12), 2353–2362.e3. 10.1016/j.cub.2020.04.015

Tsujita, M., Melchior, J. T., & Yokoyama, S. (2024). Lipoprotein Particles in Cerebrospinal Fluid. Arteriosclerosis, Thrombosis, and Vascular Biology, 44(5), 1042–1052. 10.1161/ATVBAHA.123.318284

Vera, A., Recabal, A., Saldivia, N., Stanic, K., Torrejón, M., Montecinos, H., & Caprile, T. (2015). Interaction between SCO-spondin and low density lipoproteins from embryonic cerebrospinal fluid modulates their roles in early neurogenesis. Frontiers in Neuroanatomy, 9, 72. 10.3389/fnana.2015.00072

Verdonk, C., Ajijola, O. A., & Khalsa, S. S. (2025). Toward a multidisciplinary neurobiology of interoception and mental health. Current Opinion in Neurobiology, 94, 103084. 10.1016/j.conb.2025.103084

Willnow, T. E., Hammes, A., & Eaton, S. (2007). Lipoproteins and their receptors in embryonic development: More than cholesterol clearance. Development, 134(18), 3239–3249. 10.1242/dev.004408

Wu, M.-Y., Carbo-Tano, M., Mirat, O., Lejeune, F.-X., Roussel, J., Quan, F. B., Fidelin, K., & Wyart, C. (2021). Spinal sensory neurons project onto the hindbrain to stabilize posture and enhance locomotor speed. Current Biology: CB, 31(15), 3315–3329.e5. 10.1016/j.cub.2021.05.042

Wyart, C., Carbo-Tano, M., Cantaut-Belarif, Y., Orts-Del’Immagine, A., & Böhm, U. L. (2023). Cerebrospinal fluid-contacting neurons: Multimodal cells with diverse roles in the CNS. Nature Reviews. Neuroscience, 24(9), 540–556. 10.1038/s41583-023-00723-8

Wyart, C., Del Bene, F., Warp, E., Scott, E. K., Trauner, D., Baier, H., & Isacoff, E. Y. (2009). Optogenetic dissection of a behavioural module in the vertebrate spinal cord. Nature, 461(7262), 407–410. 10.1038/nature08323

Yang, L., Rastegar, S., & Strähle, U. (2010). Regulatory interactions specifying Kolmer-Agduhr interneurons. Development (Cambridge, England), 137(16), 2713–2722. 10.1242/dev.048470

Yang, L., Wang, F., & Strähle, U. (2020). The Genetic Programs Specifying Kolmer-Agduhr Interneurons. Frontiers in Neuroscience, 14, 577879. 10.3389/fnins.2020.577879

Yue, W. W. S., Touhara, K. K., Toma, K., Duan, X., & Julius, D. (2024). Endogenous opioid signalling regulates spinal ependymal cell proliferation. Nature, 634(8033), 407–414. 10.1038/s41586-024-07889-w

Zhang, X., Jia, S., Chen, Z., Chong, Y. L., Xie, H., Feng, D., Wu, X., Song, D. Z., Roy, S., & Zhao, C. (2018). Cilia-driven cerebrospinal fluid flow directs expression of urotensin neuropeptides to straighten the vertebrate body axis. Nature Genetics, 50(12), 1666–1673. 10.1038/s41588-018-0260-3

